# Mechanistic insights into the phosphoryl transfer reaction in cyclin-dependent kinase 2: a QM/MM study

**DOI:** 10.1101/605048

**Authors:** Rodrigo Recabarren, Edison H. Osorio, Julio Caballero, Iñaki Tuñón, Jans Alzate-Morales

## Abstract

Cyclin-dependent kinase 2 (CDK2) is an important member of the CDK family exerting its most important function in the regulation of the cell cycle. It catalyzes the transfer of the gamma phosphate group from an ATP (adenosine triphosphate) molecule to a Serine/Threonine residue of a peptide substrate. Due to the importance of this enzyme, and protein kinases in general, a detailed understanding of the reaction mechanism is desired. Thus, in this work the phosphoryl transfer reaction catalyzed by CDK2 was revisited and studied by means of hybrid quantum mechanics/molecular mechanics (QM/MM) calculations. Our results show that the base-assisted mechanism is preferred over the substrate-assisted pathway, in agreement with a previous theoretical study. The base-assisted mechanism resulted to be dissociative, with a potential energy barrier of 14.3 kcal/mol, very close to the experimental derived value. An interesting feature of the mechanism is the proton transfer from Lys129 to the phosphoryl group at the second transition state, event that could be helping in neutralizing the charge on the phosphoryl group upon the absence of a second Mg^2+^ ion. Furthermore, important insights into the mechanisms in terms of bond order and charge analysis were provided. These descriptors helped to characterize the synchronicity of bond forming and breaking events, and to characterize charge transfer effects. Local interactions at the active site are key to modulate the charge distribution on the phosphoryl group and therefore alter its reactivity.

## Introduction

Cyclin-dependent kinases is a family of Serine/Threonine kinases that phosphorylate peptide substrates using adenosine triphosphate (ATP) as phosphate source with a unique function in the regulation of the cell cycle [1]. As their names state, they depend on the binding of a cyclin protein in order to be fully activated [2–4] and also on the phosphorylation of specific residues [5–9]. In particular, cyclin-dependent kinase 2 (CDK2), which can bind Cyclin E or A, needs to be phosphorylated at Thr160 [10]. CDK2 helps in the progression from G1 to S phase during the cell cycle and its malfunctioning, e.g. by mutations, has been related to different human cancers [11]. For this reason, many CDK inhibitors have been proposed, which in the majority of the cases bind in the ATP binding site. Also, and most recently, new inhibitors have been designed to tackle protein-protein interactions and allosteric sites [12]; however, the development of potent and selective inhibitors has been very challenging and in many cases disappointing, without getting through clinical trials [13]. In this context, the precise knowledge of the phosphoryl transfer mechanism in CDKs, and kinases in general, could lead to new strategies for the development of more potent and selective drugs. Moreover, CDK2 is a very interesting system to be used as a model of study within the CDK family due to the availability of many crystallographic structures and kinetic data [10,14–18].

In general, CDKs and kinases provide specific amino acids and Mg^2+^ cofactors that help to position ATP and the substrate to be phosphorylated in the correct orientation for catalysis. Also, the active site provides compensatory charges that stabilize the negative charge at the transition state (TS), lowering in this way the energy barrier of the reaction [19,20]. However, many aspects of the phosphoryl transfer mechanism are still unclear, such as the role of Mg^2+^ ions in the mechanism and which is the most favorable catalytic pathway [20]. In the former aspect, CDK2 was always thought to function efficiently with only one Mg^2+^ ion within the active site, different to what is known for other kinases, such as PKA (protein kinase A), that uses two Mg^2+^ ions as cofactors. However, recent crystallographic and kinetic data have shown that CDK2 also works more efficiently with two Mg^2+^ ions. This second Mg^2+^ ion prevents ADP release and therefore it needs to be removed from the active site after the phosphoryl transfer step is completed [21,22].

On the other hand, when it comes to the phosphoryl transfer mechanism itself, two scenarios are possible: one in which the proton from the hydroxyl group of the Ser/Thr side chain is transferred to a nearby aspartate residue (Asp127 in CDK2), route that is called base-assisted mechanism; or a second one where the proton can be transferred to one of the oxygen atoms of the γ-phosphate of the ATP molecule, pathway that is called substrate-assisted mechanism (Fig 1) [23–25]. These two mechanisms are associated to another classification of phosphoryl transfer mechanisms, named as dissociative, concerted and associative mechanisms [20]. In a dissociative mechanism, the breaking of the P-O bond with the leaving oxygen atom precedes bond formation and the reaction goes through a metaphosphate intermediate linked by two TSs. In this case, the TS is dissociative in nature or “loose”, characterized by long P-O distances with respect to the leaving and entering oxygens. In an associative mechanism, the distances of the breaking and forming P-O bonds are much shorter and the reaction goes through a pentavalent phosphorane intermediate; in this way the TS is associative or “tight”. Also, the mechanism can be concerted, which implies a single TS, where the TS may have a more dissociative or associative nature depending on the bond orders of the bonds to be broken and formed.

**Fig 1.**
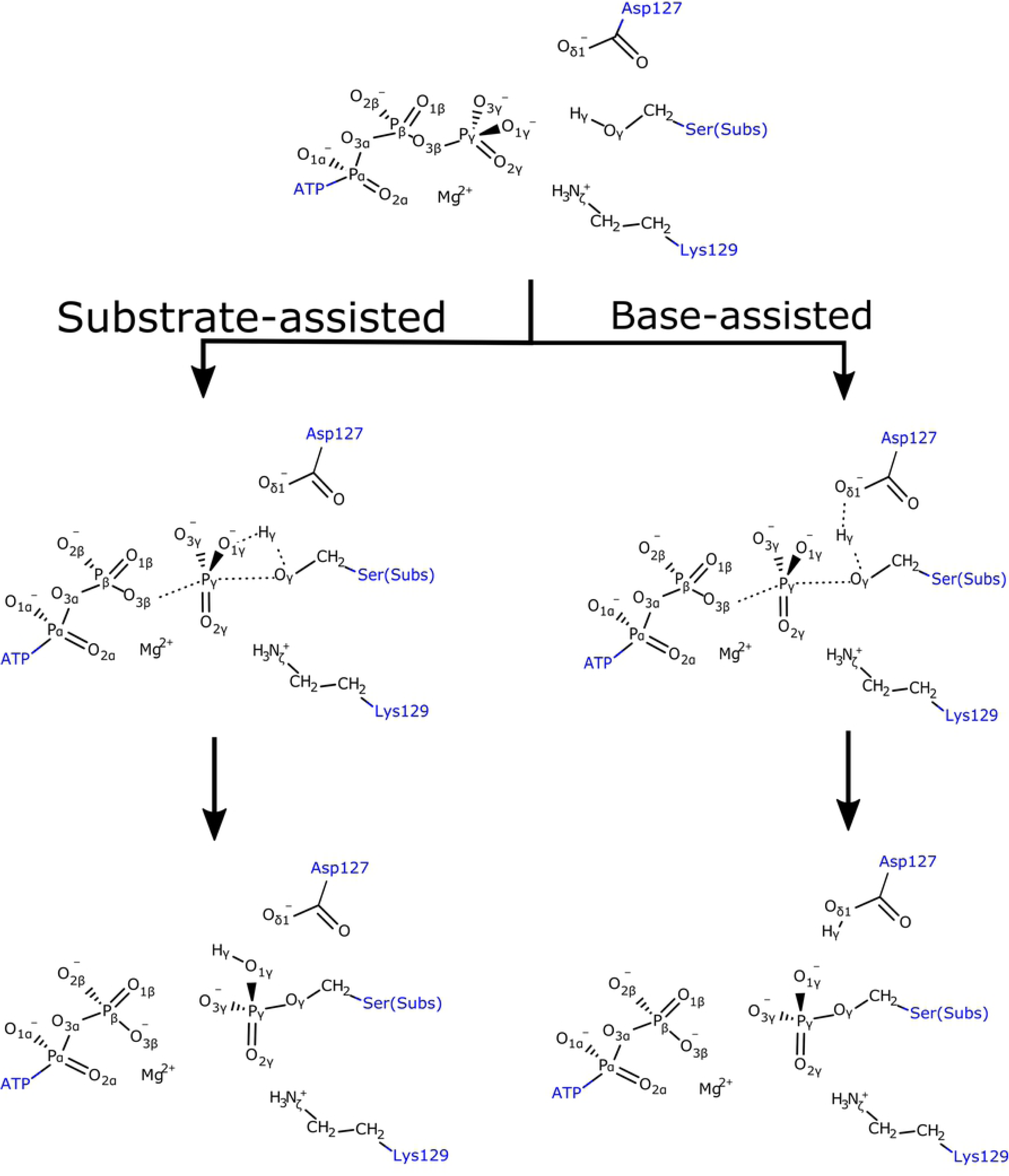
Proposed pathways for the substrate-assisted and base-assisted mechanisms. Dotted lines represent bonds that are broken or formed during the reaction.

Previous computational studies in CDK2 have not completely defined which is the most favorable route [24,26]; however, insights that the base-assisted mechanism is preferred have been given more recently [25]. The first computational study used cluster calculations at a DFT (density functional theory) level and proposed the substrate-assisted mechanism as the operating one [26]. A subsequent study carried out by the same authors, but now considering a QM/MM (quantum mechanics/molecular mechanics) partition of the system, also reaffirmed that fact, with an energy barrier for the reaction much lower than the previous one obtained (42 and 24 kcal/mol, respectively) [24]. These results proposed a structural role for Asp127, however, the residue was not included in the QM region, and therefore its participation as a catalytic base could not be ruled out. More recently, Smith *et al.* [25] by means of QM/MM calculations, found a free energy barrier of 10.8 kcal/mol for the base-assisted mechanism and an energy barrier that could be higher than 30 kcal/mol for the substrate-assisted mechanism, since the energy barrier was not estimated once they observed a monotonically increment of the energy along the reaction pathway above 30 kcal/mol. As a conclusion, they proposed the base-assisted mechanism as the most favorable one, despite the fact that they could not estimate accurately the barrier for the substrate-assisted mechanism. It is noteworthy that the calculated energy barrier was somewhat lower than the experimentally derived value (using transition state theory), which would amount to 15.3 kcal/mol (k_3_=35 s^−1^) [10]. At this point, they recall that this kinetic parameter was estimated using a different peptide substrate that showed a different amino acidic sequence. For instance, it had a Thr instead of a Ser residue to be phosphorylated. The authors proposed that this fact could explain the discrepancy. Among the limitations that this last study revealed is the small QM region used (only 48 atoms), which did not include most of the residues coordinating the Mg^2+^ cofactor, this due to the costly sampling at a DFT level. Nonetheless, it is known that this simplification could lead to overpolarization of the QM fragments, and specially in this case the coordinating metal, affecting the energetics of the system [27]. Furthermore, the free energy barrier is dependent on the selection of the reaction coordinate, which they showed can vary considerably depending on how the combination of the reactive distances is made.

Thus, in this study the goal is to revisit the phosphoryl transfer mechanism in CDK2 with one Mg^2+^ ion, allowing a more direct comparison with previous computational studies; and also to compare both mechanisms with the same QM/MM methodology, aiming at clarifying discrepancies observed in previous investigations. Besides, a detailed analysis of bond orders and atomic charges is performed in order to characterize in detail the nature of the transition states and every step of the mechanisms.

## Methods

### Protein preparation and molecular dynamics (MD) simulations

The initial coordinates of CDK2 were retrieved from the crystal structure with PDB (protein data bank) code 1QMZ [17]. This structure contains CDK2, a bound peptide with sequence HHASPRK, cyclin A3, an ATP molecule, and Mg^2+^ as cofactor. Addition of hydrogen atoms, assignation of bond orders, and correction of bond types was done with the *Protein Preparation Wizard* tool included in *Maestro* [28–30]. Three residues were restored to the crystal structure: Arg297 and Leu298 on the N-terminus of CDK2, and Glu174 on the C-terminus of cyclin A3. Protonation states at physiological pH were assigned using the *Propka* program [31]. Titratable residues were carefully analyzed by visual inspection of hydrogen networks around them. Then, a short minimization of the structure, up to a RMSD (root-mean-square deviation) convergence criterion of 0.3 Å, was performed to eliminate any steric clashes among atoms. Parameters for all residues were derived from the OPLS-2005 force field [32].

The system was immersed in a rectangular box of water molecules using the SPC model [33] and net charge neutralization was achieved by addition of two Na^+^ ions. A buffer region of 10 Å was imposed over all sides of the protein surface. Periodic boundary conditions were set over the *x*, *y* and *z* axes. The default relaxation protocol implemented in *Desmond* 2013 [34] was used. This consists in six steps in which the system is first energy minimized by a steepest descent algorithm applying and releasing restraints over heavy atoms. After that, four short MD simulations of 12 and 24 ps are performed retaining restraints, to finally carry out an unrestrained simulation. In order to study the viability of the substrate-assisted mechanism, a 3.4 ns production run was performed and the last snapshot was chosen for further QM/MM studies. On the other hand, to study the base-assisted mechanism, an extra simulation was carried out to bring together the H atom of the Ser residue to be phosphorylated with the oxygen atom of Asp127 to a geometry that may favor the proton transfer. Here, using a 5 ns MD simulation, the distance between these two atoms, and also the distance between the phosphorus atom of the γ-phosphate with the nucleophile oxygen atom of the Ser residue, were restrained with a force constant of 20 kcal/mol•Å^2^ at equilibrium distances of 1.6 and 3.4 Å, respectively. Finally, distance restraints were removed and a 2 ns run served to extract a representative structure for QM/MM calculations. It is worth noting that to all production runs a small restraint of 0.1 kcal/mol•Å^2^ was applied on the backbone atoms to keep the secondary structure of the protein stable. A 9 Å cutoff was used for the evaluation of van der Waals and electrostatic interactions, while long-range electrostatic interactions were treated with the particle mesh Ewald (PME) method [35]. Pressure was kept constant at 1 atm and temperature at 300 K using the Martyna-Tobias-Klein barostat [36] and the Nosé-Hoover chain thermostat [37], respectively. The RESPA (time-reversible reference system propagator algorithm) integrator [38] was applied with a 2 fs time step for bonded and short-range nonbonded interactions and 6 fs for long-range electrostatics.

### QM/MM calculations

The last snapshot from MD simulations was used for setting QM/MM calculations. The QM region consisted in the triphosphate moiety of the ATP molecule, side chains of residues Asp145, Asn132, Lys129, Asp127, Ser5 (residue to be phosphorylated), the Mg^2+^ ion, and a coordinating water molecule (Fig 2). All QM/MM cuts were done between the Cβ and Cα atoms and valences were saturated with hydrogen atoms (link-atom approach), resulting in a total of 61 atoms in the QM region. Transition state optimizations were done with a micro-macro iterative procedure [39]. After that, the intrinsic reaction coordinate (IRC) path was traced down to reactant and product states, with full optimization of them. To speed-up the calculations, only the atoms within a radius of 25 Å from the ATP molecule were allowed to move, while the rest of MM atoms were kept frozen. Treatment of QM atoms was done at the B3LYP/6-31+G* [40] level of theory, similar to what has been used in previous investigations [25], while the classical OPLS-AA force field [41] was used for MM atoms, as implemented in the *fdynamo* library [42]. Electronic properties such as charges and Wiberg bond orders were computed for each point along the IRC path using the natural bond orbital method (NBO) [43,44]. All calculations were done with *fdynamo* [42] interfaced with the *Gaussian 09* program [45].

**Fig 2.**
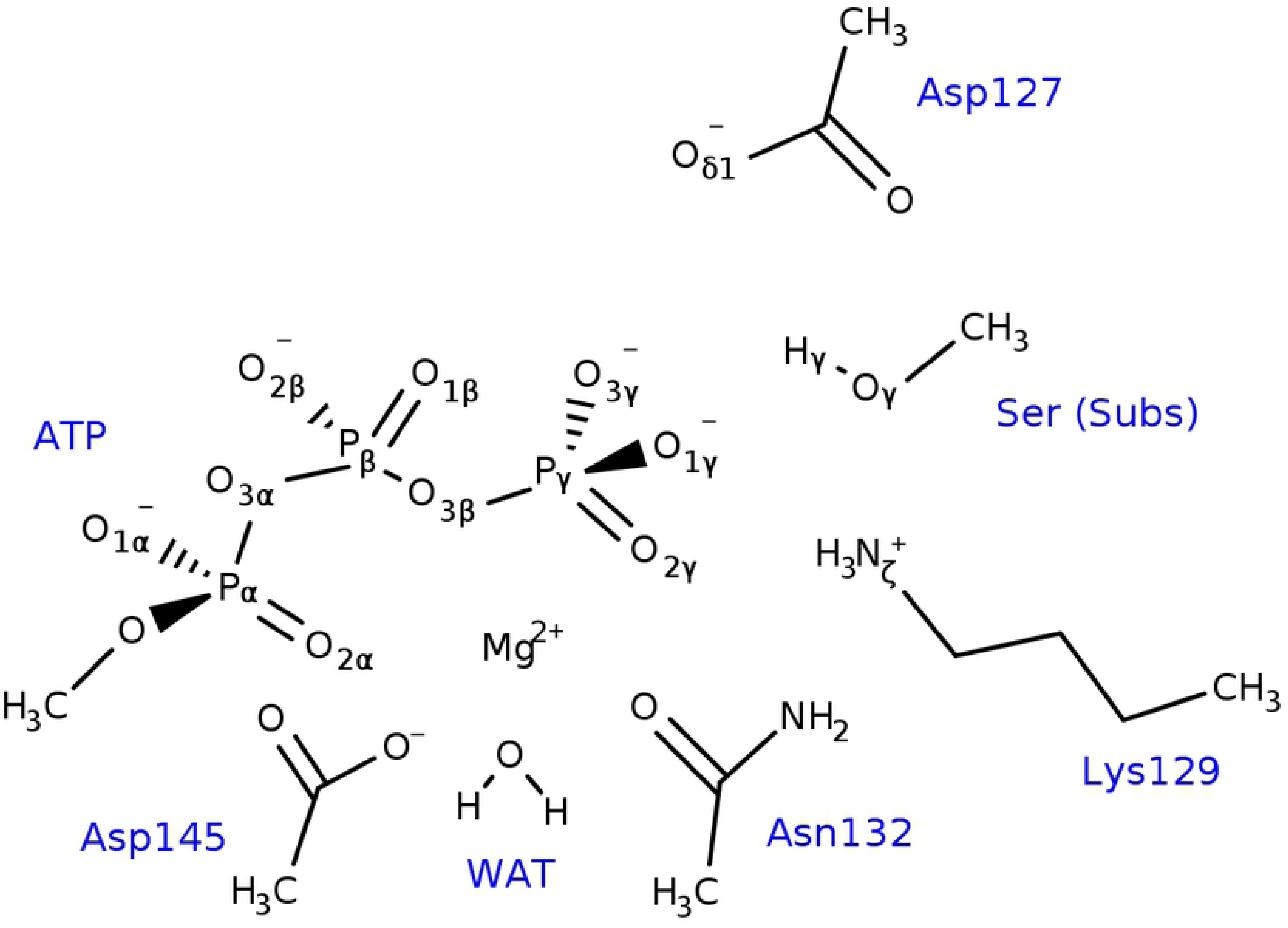
Atoms included in the QM region for QM/MM calculations. The valences of the atoms lying at the QM/MM boundary were completed with hydrogen atoms.

## Results and discussion

### Potential energy barriers

The potential energy barriers for the substrate-assisted and base-assisted mechanisms were estimated and their values are shown in Fig 3. It is possible to see how the substrate-assisted route exhibits a much higher activation energy (50.5 kcal/mol) compared to the base-assisted pathway (14.3 kcal/mol). Also, for the former mechanism, only one TS was found, making this route concerted. In the other case, two TSs were identified, which are flanked by an intermediate structure; however, the energy difference between the first transition state (TS1) and the intermediate structure (Int) is of only 1.1 kcal/mol. The energy difference between Int and TS2 is much more marked (7.2 kcal/mol), where the product state is finally reached with a reaction energy of 2.5 kcal/mol, which is very similar to the substrate-assisted mechanism (2.3 kcal/mol). Here, and as it was mentioned, the energy difference between TS1 and Int is very small and therefore the stepwise/concerted nature of the mechanism could change when moving from potential energy surfaces to free energy surfaces. Ongoing work in our lab is being conducted to clarify this point.

**Fig 3.**
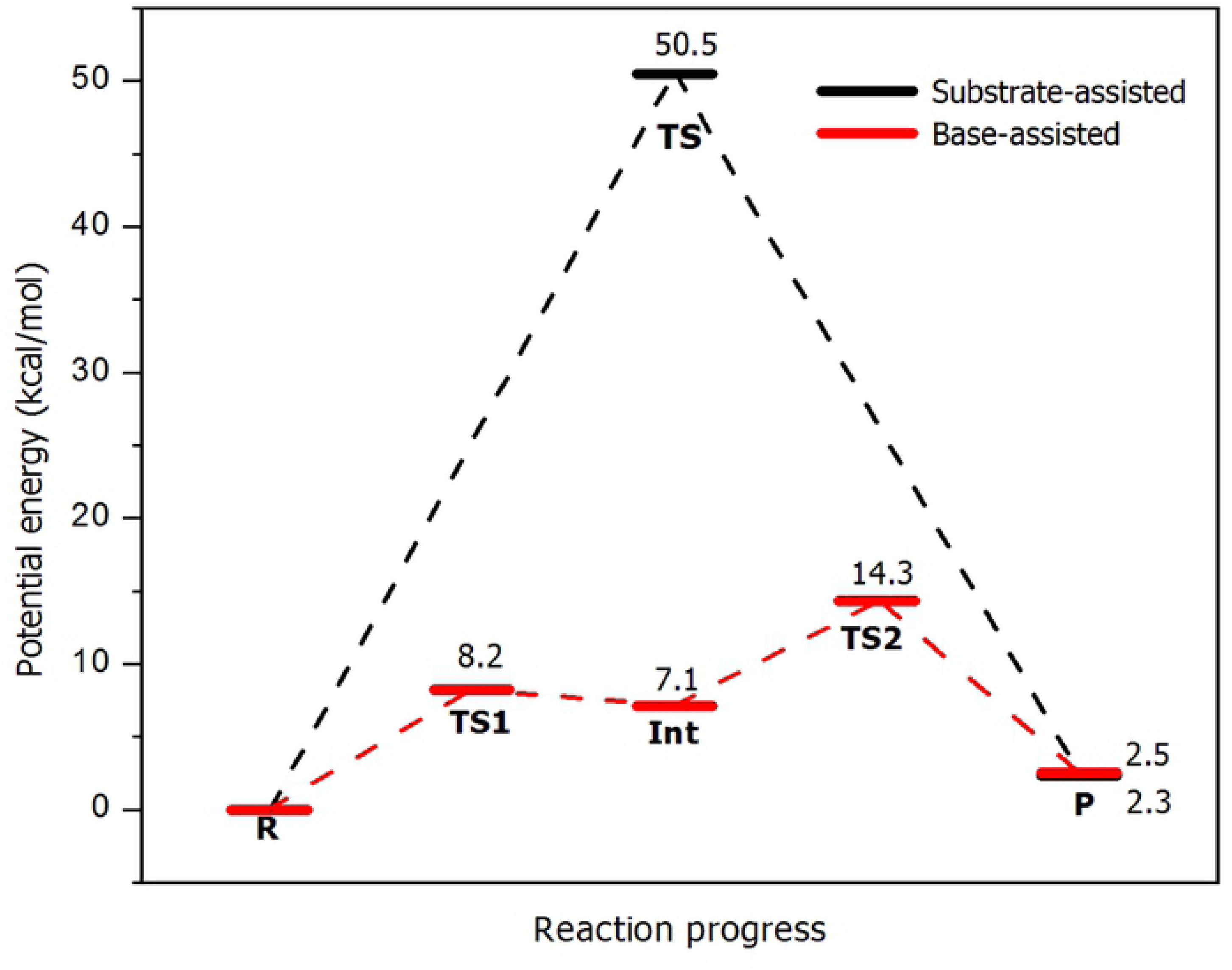
Potential energy profiles for both, substrate-assisted (black) and base-assisted (red) mechanisms. The numbers show the relative energy for each stationary point with respect to reactants.

According to the calculated energy barriers, the base-assisted mechanism is more favorable than the substrate-assisted pathway, what is in agreement with the last computational study carried out by Smith *et al.* [25]. It is worth noting that the calculated activation energy for the base-assisted mechanism (14.3 kcal/mol) agrees very well with the experimental derived value of 15.3 kcal/mol, making this mechanism the most probable one. Regarding the substrate-assisted mechanism, our value for the activation energy (50.5 kcal/mol) could be overestimated, since other studies in CDK2 and PKA have shown that energy barriers for this pathway are within a range of 20-40 kcal/mol [24,46–48]. However, there are also computational studies in PKA that have shown activation energies higher than 45 kcal/mol [47,49], closer to the value obtained in our study. On the other hand, an additional test was performed carrying out single point calculations on the optimized geometries using the M06-2X functional [50], but a very similar energy barrier was obtained. In this context, a plausible explanation for the high-energy barrier is the formation of a highly strained four-membered ring (P_γ_-O_1γ_-H_γ_-O_γ_, see Fig 1) at the TS, effect that has also been proposed in other studies [46,49].

### Structural analysis of reactants, TSs and products

The reactants state obtained for both reaction mechanisms was different since the conformation used was extracted from different MD simulations. In the case of the substrate-assisted mechanism, the Ser residue from the peptide substrate is establishing a hydrogen bond contact with the oxygen O_1γ_ of the γ-phosphate (see Figs 1 and 4A), thereby allowing the proton transfer reaction to take place to this atom. In the case of the base-assisted mechanism, an extra simulation was performed to generate a hydrogen bond between the hydroxyl group of the substrate Ser residue and residue Asp127, which showed to be stable during the MD simulation (S1 Fig.). The optimized geometries of the reactants states for both reaction mechanisms are shown in Figs 4A and 5A, respectively. In the case of the substrate-assisted mechanism, the initial distance between the P_γ_ atom and the entering oxygen O_γ_ is 3.98 Å, slightly longer than the crystallographic distance (3.68 Å, Table 1). The TS in the substrate-assisted mechanism (Fig 4B) shows that the phosphoryl group is midway between the donor O_3β_ atom and the acceptor O_γ_ atom from the Ser residue, but pushed towards the entering oxygen atom (O_3β_-P_γ_=3.04 Å vs P_γ_-O_γ_=2.10 Å, Table 1), resembling an almost planar geometry. Using Pauling’s formula to assess the degree of associativity for a phosphoryl transfer reaction [51], D(n)=D(1) – 0.60log(n), where D(1) is the P-O distance for a single bond (1.73 Å), and with D(n) being the average between the two P-O distances (2.57 Å), the fractional bond number (n), which gives a measure of the associativity of the mechanism, can be calculated. The value obtained for n amounts to 0.039, which means the TS is 3.9 % associative or 96.1 % dissociative. This result shows that though the mechanism is concerted, the TS is quite dissociative or loose, which agrees with the rather large distance between the leaving and entering oxygen atoms at the TS (4.91 Å). On the other hand, at the TS, the proton transfer reaction has also started but is not totally completed, showing the H atom between both donor and acceptor oxygen atoms (Fig 4B). Fig 4C shows the product state, where it is possible to observe how the phosphoryl transfer reaction has been completed, with the characteristic that the transferred proton is forming a hydrogen bond with one of the oxygen atoms of the β-phosphate (O_3β_) of the newly formed ADP molecule. The Lys129 and Asp127 residues help in maintaining the in-line configuration necessary for the phosphoryl transfer reaction to take place and, especially, Lys129 stabilizes the transferred phosphoryl group at the TS through hydrogen bonding interactions. It is also possible to observe that the octahedral coordination around the Mg^2+^ ion is preserved during and after the reaction (Table 1), in agreement with previous computational studies [24,25].

**Fig 4.**
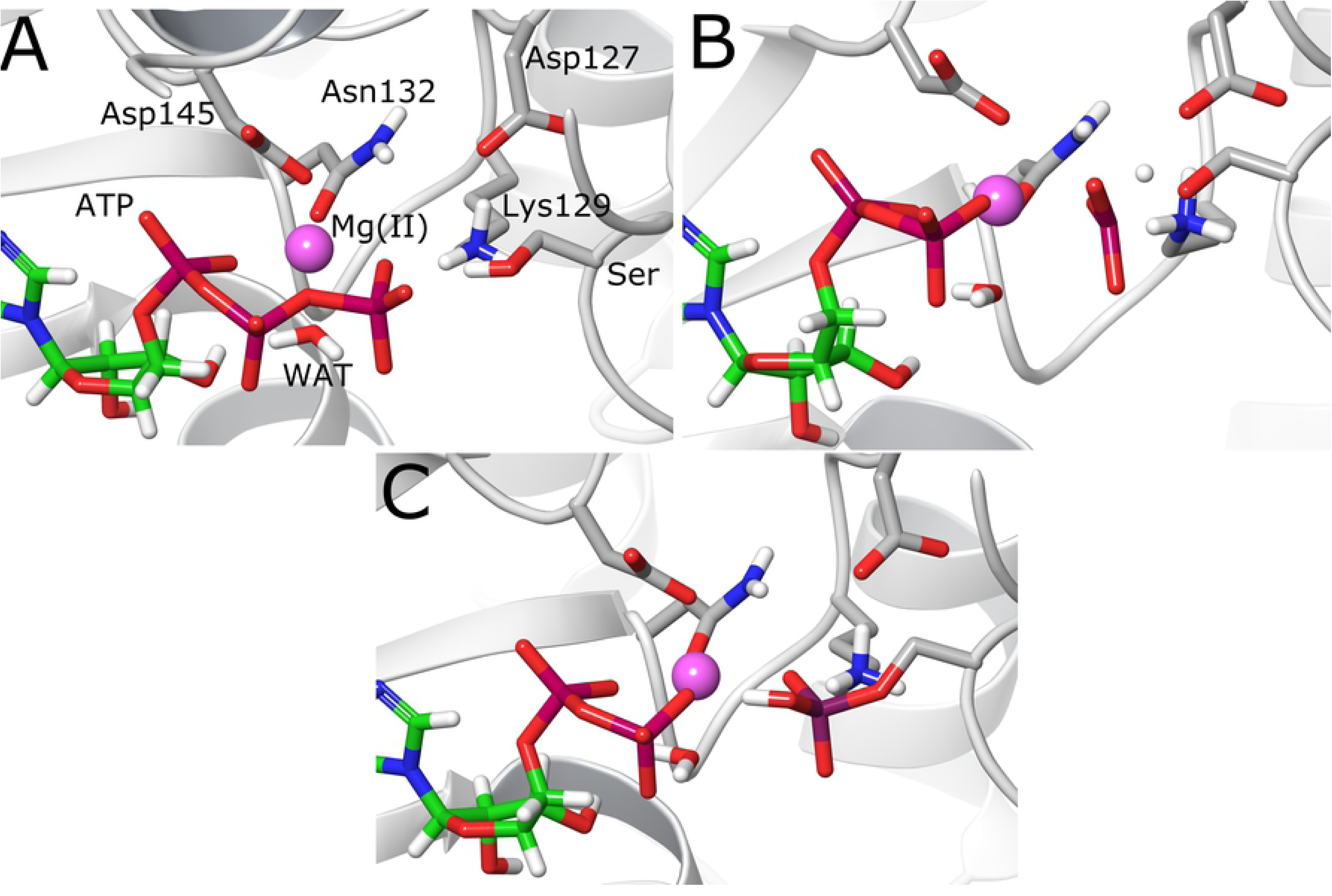
Molecular structures of the stationary points in the substrate-assisted mechanism. (A) Reactant state. (B) Transition state. (C) Product state.

**Table 1.** Measured distances between key atoms that participate during catalysis and those that complete the octahedral coordination around the Mg^2+^ ion.

| Distance (Å) | Exp.<sup>a</sup> | Substrate-assisted |  |  | Base-assisted |  |  |  |  |
| --- | --- | --- | --- | --- | --- | --- | --- | --- | --- |
|  |  | Reac | TS | Prod | Reac | TS1 | Int | TS2 | Prod |
| O<sub>γ</sub>(Ser)-P<sub>γ</sub> | 3.68 | 3.98 | 2.10 | 1.65 | 3.19 | 2.47 | 2.12 | 1.83 | 1.70 |
| O<sub>3β</sub>-P<sub>γ</sub> | 1.60 | 1.76 | 3.04 | 3.39 | 1.82 | 2.57 | 2.95 | 3.46 | 3.66 |
| O<sub>γ</sub>(Ser)-H<sub>γ</sub>(Ser) |  | 0.99 | 1.24 | 3.38 | 0.98 | 0.99 | 1.00 | 1.19 | 1.84 |
| O<sub>δ1</sub>(Asp127)-H<sub>γ</sub>(Ser) |  | 4.70 | 4.79 | 5.65 | 1.75 | 1.76 | 1.72 | 1.30 | 1.00 |
| O<sub>1γ</sub>-H<sub>γ</sub>(Ser) |  | 1.83 | 1.22 | 1.00 | 3.36 | 3.06 | 2.96 | 3.11 | 3.38 |
| Mg-O<sub>2γ</sub> | 2.25 | 2.04 | 2.39 | 2.39 | 2.06 | 2.23 | 2.39 | 2.98 | 3.18 |
| Mg-O<sub>3β</sub> | 2.12 | 2.33 | 1.98 | 2.02 | 2.18 | 2.02 | 1.99 | 1.98 | 1.97 |
| Mg-O<sub>2α</sub> | 2.07 | 2.03 | 2.11 | 2.19 | 2.05 | 2.07 | 2.09 | 2.02 | 2.03 |
| Mg-O<sub>WAT</sub> | 2.14 | 2.27 | 2.20 | 2.17 | 2.17 | 2.18 | 2.19 | 2.17 | 2.17 |
| Mg-O(Asn132) | 2.00 | 2.14 | 2.45 | 2.58 | 2.08 | 2.18 | 2.22 | 2.18 | 2.17 |
| Mg-O(Asp145) | 1.91 | 2.02 | 2.01 | 2.01 | 2.03 | 2.02 | 2.01 | 1.98 | 1.98 |
| N<sub>ζ</sub>(Lys129)-O<sub>2γ</sub> | 3.58 | 2.83 | 2.82 | 2.73 | 2.72 | 2.69 | 2.66 | 2.52 | 2.65 |
| N<sub>ζ</sub>(Lys129)-H<sub>1ζ</sub>(Lys129) |  | 1.06 | 1.04 | 1.05 | 1.06 | 1.05 | 1.06 | 1.40 | 1.64 |
| O<sub>2γ</sub>-H<sub>1ζ</sub>(Lys129) |  | 1.80 | 1.84 | 1.73 | 1.70 | 1.65 | 1.61 | 1.12 | 1.03 |

<sup>a</sup>Experimental values are derived from the crystal structure with PDB code 1QMZ [17]

In the case of the base-assisted mechanism, Fig 5B shows how in the first TS (TS1) the phosphoryl group has a planar geometry with almost equal distances between P_γ_ and the leaving and entering oxygen atoms (2.57 and 2.47 Å, respectively). The distance between those two oxygen atoms is 5.03 Å, revealing the high dissociative character of the mechanism, and with a fractional bond number of 0.048, i.e., 95.2 % dissociative. Very close in energy is the intermediate structure (Int, Fig 5C), which also resembles a metaphosphate geometry but with a shorter distance to the entering oxygen atom (2.12 Å, Table 1). At this point, the distance of the oxygen atom O_2γ_ coordinating the Mg^2+^ ion has increased from 2.06 Å to 2.39 Å. The reaction proceeds and a second transition state (TS2) is reached, where the P_γ_-O_γ_ bond is almost formed (1.83 Å). At this stage, the proton transfer to Asp127 has been initiated and the proton is midway between the donor (O_γ_) and acceptor (O_δ1_) oxygen atoms. An interesting feature of this structure is the spontaneous transfer of a proton from Lys129 to the O_2γ_ atom of the ATP phosphoryl group. This proton transfer reaction is almost complete at this transition state, with the proton lying between the donor nitrogen and acceptor oxygen atoms (Fig 5D). Here, it is very probable that neutralization of Asp127 plays a role in lowering the pK_a_ of Lys129, favoring in this way the proton transfer. Finally, the system reaches the product state (Fig 5E), which features the phosphorylated Ser residue with a distance between P_γ_ to the O_3β_ oxygen of 3.66 Å. Asp127 is protonated forming a hydrogen bond with the oxygen atom of the Ser residue. The phosphorylated Ser residue is now protonated at the O_2γ_ oxygen and forming a hydrogen bond with the nitrogen atom of Lys129. The O_2γ_ oxygen has partially lost its coordination with the Mg^2+^ ion, since the distance to it has increased to 3.18 Å. All the rest of the O-Mg^2+^ distances are in general conserved in values that do not go beyond 2.2 Å (Table 1). Finally, it is expected that the usual protonation states of Asp127 and Lys129 may be easily recovered when the products leave the active site.

**Fig 5.**
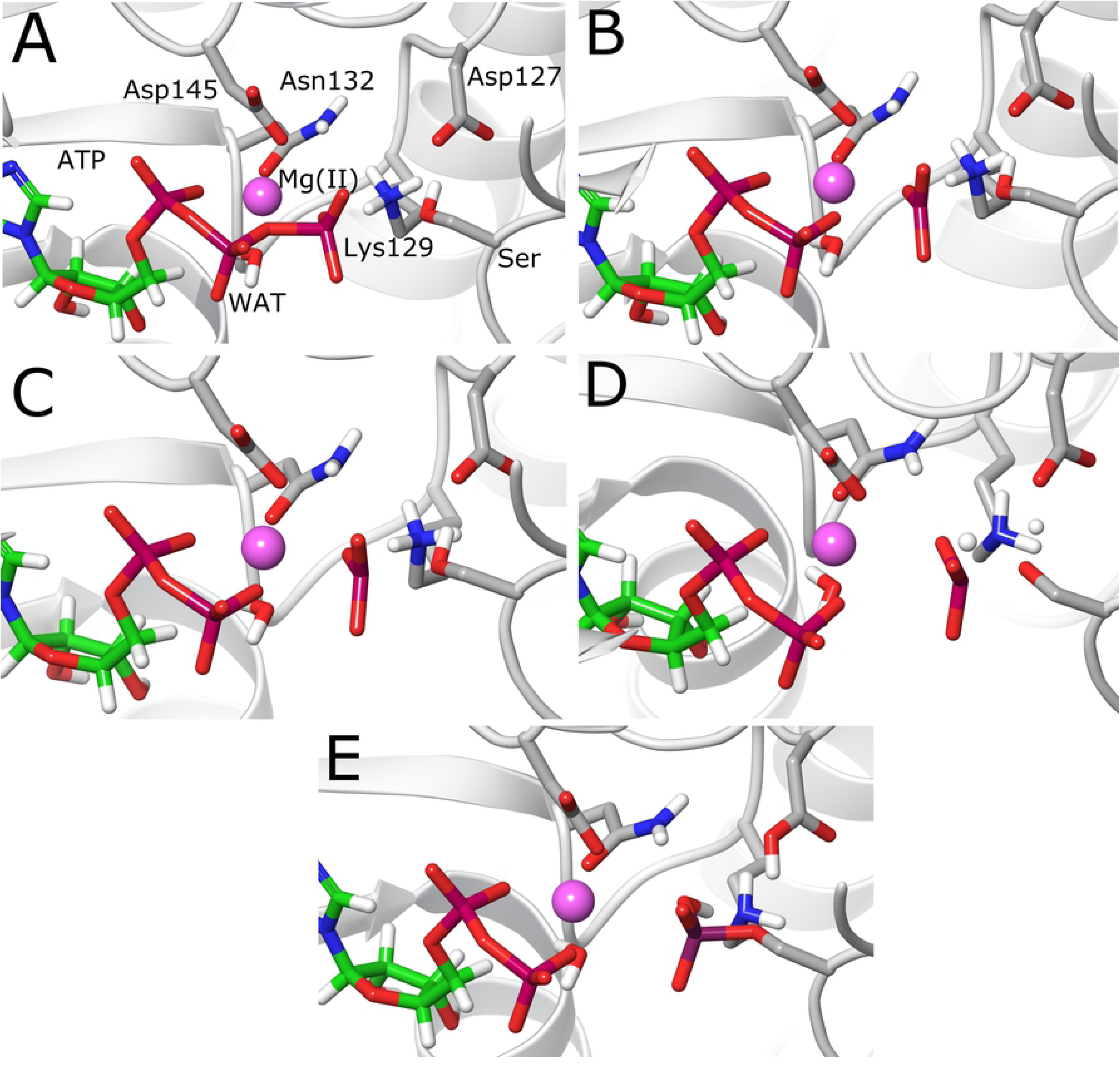
Molecular structures of the stationary points in the base-assisted mechanism. (A) Reactant state. (B) Transition state TS1. (C) Intermediate state Int. (D) Transition state TS2. (E) Product state.

These results show that while the substrate-assisted mechanism exhibits a concerted character, the base-assisted mechanism is dissociative. Asp127 serves as a base accepting the proton from the substrate Ser residue late in the progress of the reaction, what also agrees with the last computational study in CDK2 [25] and with QM/MM calculations in PKA [46]. It is interesting that in both mechanisms the TSs exhibit a dissociative character, though the substrate-assisted mechanism has been related to more associative mechanisms [24,46]. The possible cause for this could be the absence of the second Mg^2+^ in our simulations, which would reduce the repulsion between the reactive fragments. On the other hand, one of the new features found in the base-assisted mechanism is the proton transfer from Lys129 to the transferred phosphoryl group. This result would suggest that Lys129 not only exerts its structural function bridging closer the γ-phosphate and the Ser residue, but could also help to stabilize the negative charge created on the phosphoryl group at the TS by a proton transfer reaction. A similar charge stabilizing event was also observed by QM/MM calculations in Glucokinase [52], where Lys169 was identified as an acid catalyst protonating one of the oxygen atoms of the γ-phosphate in the ATP molecule. The great structural similarity among protein kinases allows for some comparison of the mechanisms since both enzymes share the common residues in the active site that are key for enzyme catalysis. In the case of CDK2, as mentioned previously, it is now postulated that two Mg^2+^ ions are required for optimal catalysis [21,22]. Despite that this structural effect is out of the scope of the present investigation, it is plausible to think that in the absence of a second Mg^2+^ ion, which would neutralize the negative charge in the active site, a proton transfer reaction from Lys129 may help to mimic its charge stabilizing function.

### Synchronicity studied by bond order analysis

As previously described, single point calculations using the NBO method [53] on each point along the reaction paths were performed in order to calculate Wiberg bond orders. Bond orders are a direct measurement of the formation or breakage’s degree of a bond and therefore are a good property to describe the synchronicity of bond breaking and bond forming events. In order to get clearer insights into these events, it has been postulated the use of bond order derivatives, as a useful qualitative property to describe synchronicity and electronic changes in chemical reactions [54,55]. These will show positive values for bond formation or strengthening, while negative values are expected for bond breaking or weakening. Figs 6A and 6B show the evolution of the bond orders for both mechanisms along the reaction coordinate, while Figs 6C and 6D show bond order derivatives. It is readily seen from Fig 6A that bond orders follow the expected behavior for the substrate-assisted reaction: the bond O_3β_-P_γ_ begins to break gradually while the rest of the bonds are maintained almost unaltered until this bond is completely broken; then, the bond P_γ_-O_γ_ begins to form. This supports the fact that P_γ_ at the TS structure shows a longer distance with respect to the leaving oxygen (O_3β_) than with the entering oxygen (O_γ_). At the TS, the curves representing the bonds O_γ_-H_γ_, O_1γ_-H_γ_ and P_γ_-O_γ_ cross with a value for the bond order of approximately 0.6 (S1 Table), showing that at the TS the formation and cleavage of these bonds has advanced halfway. This also shows that there is a marked synchronicity in the formation and dissociation of these bonds. However, this is better illustrated in Fig 6C, where bond order derivatives show an initial negative sign for the O_3β_-P_γ_ bond, depicting its gradual breaking, and how the peaks for the other three bonds are located exactly at the TS, with the positive peaks representing the formation of the O_1γ_-H_γ_ and P_γ_-O_γ_ bonds, and a negative peak representing the breaking of the O_γ_-H_γ_ bond. Thus, these results show that though the reaction is concerted, there is a high degree of asynchronicity with respect to the breaking of the O_3β_-P_γ_ bond, while the rest of the events occur in a synchronous fashion. Furthermore, it is seen how the peak representing the formation of the P_γ_-O_γ_ bond is quite broad in direction to the product state, showing how the strengthening of this bond is gradual. Finally, the last negative peak in the O_1γ_-H_γ_ bond represents some elongation that this bond undergoes when start forming a hydrogen bond with the O_3β_ oxygen of ADP.

**Fig 6.**
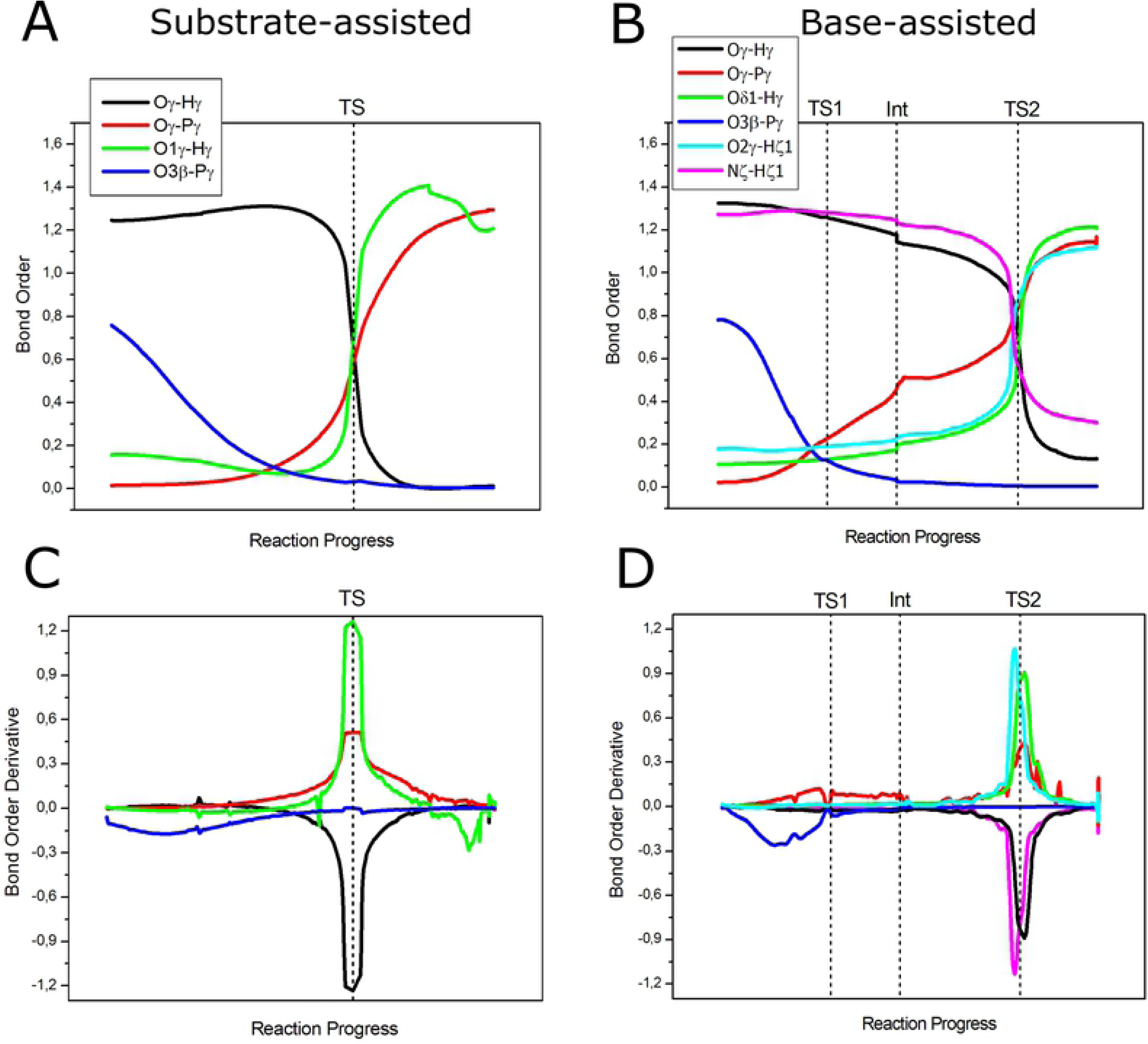
Wiberg bond orders and bond order derivatives of the bonds that are broken or formed during the reaction for both mechanisms. (A) Bond orders for the substrate-assisted mechanism. (B) Bond orders for the base-assisted mechanism. (C) Bond order derivatives for the substrate-assisted mechanism. (D) Bond order derivatives for the base-assisted mechanism. Dotted lines show the position of the stationary points on the reaction coordinate.

In the case of the base-assisted mechanism, bond breaking and forming events occur in a similar way, beginning with the breaking of the O_3β_-P_γ_ bond (Fig 6B) until TS1 is reached. At this stage, the bond P_γ_-O_γ_ has just started to form and the reaction proceeds with the strengthening of this bond until the intermediate structure is formed. At this point, this bond is almost half-formed, i.e., bond order of 0.46 (S2 Table), while the bond O_3β_-P_γ_ is completely broken. Following the reaction coordinate, it is possible to observe small variations in the bond orders for the rest of the bonds due to structural rearrangements prior the completion of the phosphoryl and proton transfer reactions. As was discussed previously, at TS2 the phosphoryl transfer is almost complete, with a bond order value of 0.83 for the P_γ_-O_γ_ bond and with intermediate values for the rest of the bonds that participate in both proton transfers (S2 Table). Fig 6D shows clearly the degree of synchronicity of bond forming and breaking events. As expected, the first negative peak represents the breaking of the O_3β_-P_γ_ bond, followed by positive values representing some degree of formation of the P_γ_-O_γ_ bond. In this case, bond formation with O_γ_ occurs markedly before reaching TS2, where the proton transfer reactions take place. It is also observed that the proton transfer from Lys129 to the phosphoryl group occurs in a slightly asynchronous way with respect to the proton transfer to Asp127 and the strengthening of the P_γ_-O_γ_ bond, i.e., though the peaks are very close, they do not have their maxima at the same position on the reaction coordinate. The positive and negative peaks representing the proton transfer from Lys129 to the phosphoryl group are located slightly before the peaks representing the proton transfer to Asp127 and the completion of the P_γ_-O_γ_ bond. This is also revealed by the bond order values of the bonds that are formed at TS2 (O_2γ_-H_ζ1_=0.87 vs O_δ1_-H_γ_=0.62, S2 Table).

### Charge analysis

In order to gain a deeper insight into the differences between the two explored reaction paths and to evaluate possible charge transfer effects, NPA (natural population analysis) charges [56] were calculated and are plotted in Fig 7. Figs 7A and 7B show the evolution of the charge on the P_γ_ atom along the reaction coordinate. It is observed that the trend of the atomic charge on this atom follows the same behavior in both mechanisms: the charge decreases (more negative values) as the mechanism approaches the TS region and increases its value until the product state. In the case of the substrate-assisted mechanism, the initial value is 2.55 (charges are presented in a.u. in this section) which decreases to a value of 2.49 before reaching the TS. At the TS, it takes a value of 2.52, from where increases until reaching a value of 2.61 at the product state (S3 Table). Thus, though there are changes on the charge of this atom, these fluctuations are not large. This event is accompanied by a shallow increase in the charge of the Mg^2+^ ion (Fig 7E), showing that this ion loses some electron density as the coordination with O_2γ_ is weakened. It is also seen that the charge on the non-bridging oxygen atoms increases (more positive) slightly and then decreases before reaching the TS, especially for O_1γ_ and O_3γ_ (Fig 7C). This behavior can be explained since the breaking of the O_3β_-P_γ_ bond withdraws electron density from the P_γ_ atom and charge transfer from the non-bridging oxygen atoms would take place to compensate for the created electron deficiency in P_γ_. On the other hand, it is also observed that the non-bridging oxygens O_2γ_ and O_3γ_ possess a similar atomic charge, but with O_2γ_ bearing a slightly more negative charge, since it is expected to be more polarized by the coordination with the Mg^2+^ ion. On the other hand, the oxygen atom O_1γ_ has a more positive charge, most probably since it is only stabilized by one hydrogen bond with the hydroxyl group of the Ser residue (Fig 8A). Otherwise, O_3γ_ is initially stabilized by three hydrogen bonds established with three water molecules, providing a larger stabilization for the charge developed on this atom (Fig 8A). However, the hydrogen bond that is initially formed with the water molecule that coordinates Mg^2+^ is lost before reaching the TS. It is worth to note that variations in the atomic charges are more pronounced on the oxygen atoms compared to the changes on the P_γ_ atom and the Mg^2+^ ion, and therefore the changes on the phosphoryl group (PO_3_^−^) charge will be determined by the charge fluctuations on the non-bridging oxygens. Before reaching the TS, the charges on O_2γ_ and O_3γ_ are the same, since at this stage the phosphoryl group resembles a metaphosphate-like structure, this is, the bond order O_3β_-P_γ_ is close to zero and the new P_γ_-O_γ_ bond has not yet been formed.

**Fig 7.**
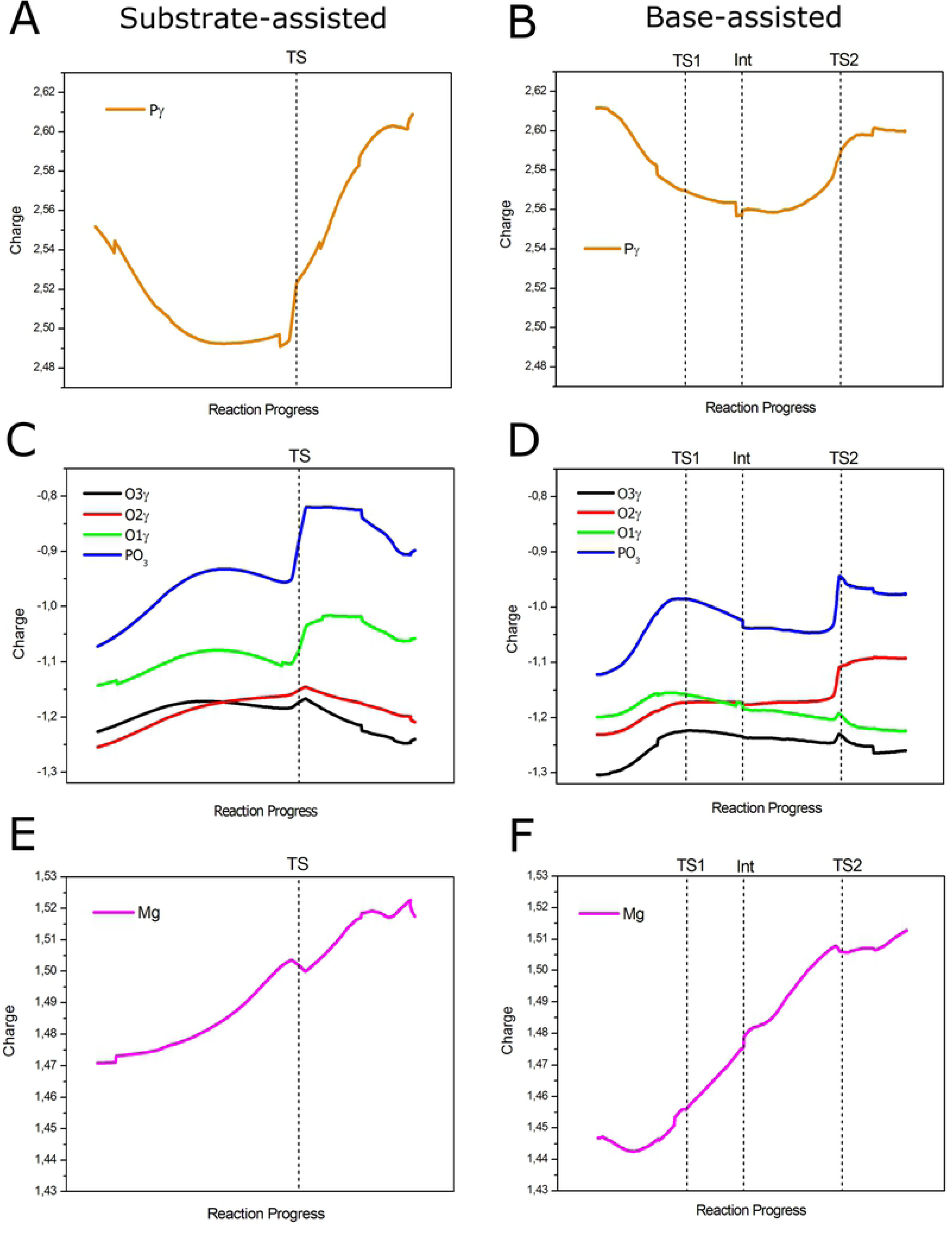
Evolution of NPA charges for the most relevant atoms in the reaction. (A, C and E) Charges for P_γ_, PO_3_ and Mg^2+^ in the substrate-assisted mechanism, respectively. (B, D and F) Charges for P_γ_, PO_3_ and Mg^2+^ in the base-assisted mechanism, respectively. Dotted lines show the position of the stationary points on the reaction coordinate.

**Fig 8.**
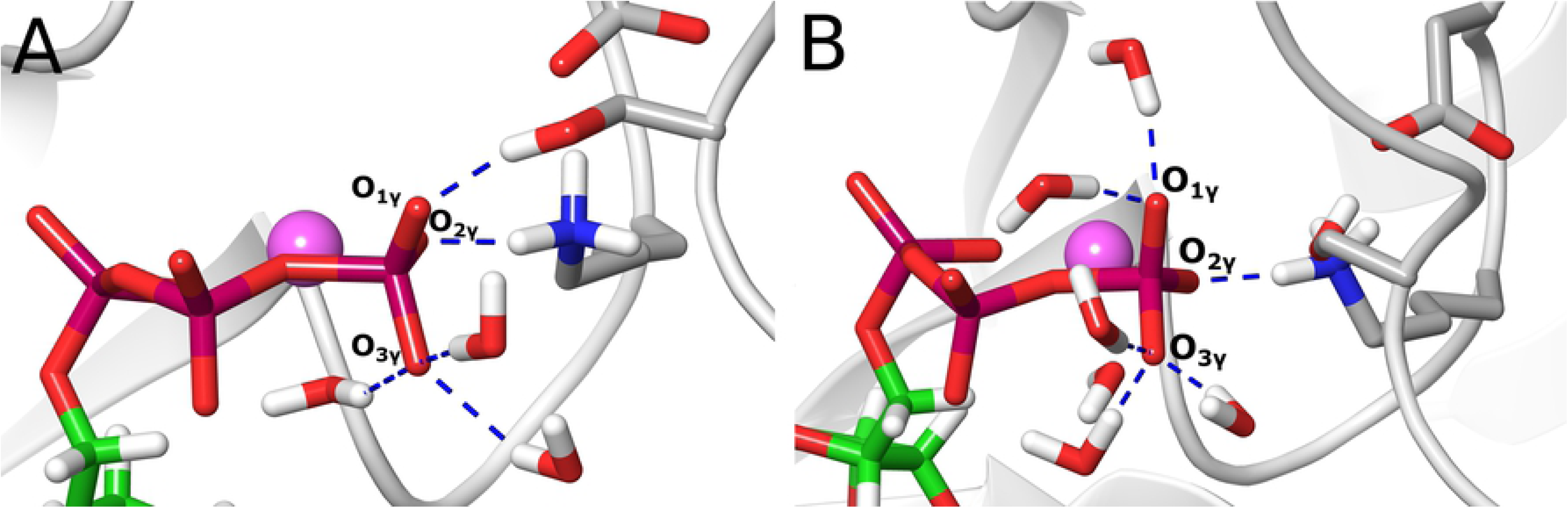
Reactants structures showing hydrogen bonding interactions (dashed blue lines) that stabilize the γ-phosphate of the ATP molecule. (A) Substrate-assisted mechanism. (B) Base-assisted mechanism.

With these observations at hand, it becomes clear that the effect that the protein environment exerts on the charge distribution of the phosphoryl group is very important and that charge stabilization depends on specific interactions at the active site, especially on hydrogen bonding interactions. At the TS, the charge on the phosphoryl group increases (more positive) sharply due to the proton transfer to the O_1γ_ atom. After the TS, the charge on the phosphoryl group remains stable and decreases before reaching the product state, accounting for the formation of the P_γ_-O_γ_ bond. The charges on O_2γ_ and O_3γ_ also become more negative due to the formation of the new bond, but now the charge on O_2γ_ becomes slightly more positive than O_3γ_, most probably since the distance with Mg^2+^ has increased and hydrogen bond stabilization on O_3γ_ is enhanced. It is also observed that the charge on Mg^2+^ increases monotonically (Fig 7E), accounting for the continuing weakening of its coordination with O_2γ_, until the formation of the product state; except for a small fluctuation at the TS, due to the change in the charge distribution of the phosphoryl group upon proton transfer.

In the case of the base-assisted mechanism, the P_γ_ atom also decreases its charge (less positive) as approaches the TS (Fig 7B). The initial value is 2.61 and then decreases to 2.57 at TS1, and it diminishes slightly more at the intermediate structure with a value of 2.56 (S3 Table). At TS2 the value has increased to 2.59, with a final value at the product state of 2.60. Interestingly, compared to the substrate-assisted mechanism, the difference in the charge value between the reactant state and the lowest value along the reaction coordinate is the same (0.06); however, the magnitude of the charge on this atom is more positive in the second path, showing that the P_γ_ atom in the base-assisted mechanism is more reactive to the attack of the entering oxygen atom, since its electrophilicity is enhanced. This effect could also help to explain the lower energy barrier for the base-assisted mechanism. As with the previous analysis, the charge on the Mg^2+^ ion smoothly increases along the reaction coordinate (Fig 7F), but in this case the difference between the charges, at the reactant and product state, is larger (0.07) when compared to the substrate-assisted mechanism (0.04). On the other hand, it is also seen that the charge on the non-bridging oxygens is slightly more negative compared to the substrate-assisted mechanism, what also makes the charge on the phosphoryl group more negative (Fig 7D).

Specifically, the charge on O_3γ_ is notoriously more negative when compared to O_1γ_ and O_2γ_. This is explained since O_3γ_ is stabilized by three hydrogen bonds from three water molecules (Fig 8B), what brings increased charge stabilization for that atom. In the case of O_1γ_, upon the absence of a hydrogen bond with the Ser residue, two hydrogen bonds with two water molecules are formed instead (Fig 8B); this results in a charge slightly more positive than for the O_2γ_ atom, which coordinates to the Mg^2+^ ion. Hence, the charge on the phosphoryl group follows the trend of the non-bridging oxygen atoms; this is, the charge increases (more positive) until reaching TS1, where it takes a value of −0.98. As discussed previously, this effect is due to the dissociation of the O_3β_-P_γ_ bond, event that withdraws electron density from the phosphoryl group. After that, the charge begins to gradually decrease until reaching the intermediate structure with a charge of −1.03. This is due to the fact that some degree of formation of the P_γ_-O_γ_ bond is already observed at this stage (see Fig 6D and its discussion), what generates a gradual recovery of the negative charge on the phosphoryl group. From this point, the charge value is practically maintained until just before reaching TS2, what agrees with the little formation of the P_γ_-O_γ_ bond between Int and TS2. Then, the charge increases to −0.94 due to the protonation of O_2γ_ by Lys129, and then diminishes slightly to a value of −0.97 at the product state, due to the completion of the P_γ_-O_γ_ bond. It is interesting to note that the charge on the phosphoryl group at TS1 and Int is close to the charge expected in a dissociative mechanism: −1 for a metaphosphate species [20]. The more negative charge on the phosphoryl group in the base-assisted mechanism correlates well with a larger change in the charge on Mg^2+^ (Fig 7F), which donates electron density to the phosphoryl group upon dissociation of the coordinating bond with O_2γ_.

In this way, our results highlight the importance of local interactions in the modulation of charge on the phosphoryl group. Furthermore, it was shown how in a reaction with high dissociative character the charge on the phosphoryl group becomes more positive when it reaches a metaphosphate species, an issue that was not clearly established in the previous literature [20]. On the contrary, for an associative mechanism, a negative charge buildup at the TS is expected, a fact that has been previously characterized computationally [57]. Our results support experimental observations using ^19^F NMR on MgF_3_^−^ transition state analogues, which have been able to describe how the direct microenvironment in near transition-state conformations alter the charge distribution of the TS mimic, i.e., by local electrostatic and hydrogen bonding interactions [58]. This is exactly what our results show for the phosphoryl group in both mechanisms: the charge distribution is finely modulated by local interactions with the Mg^2+^ ion, Lys129, the substrate Ser residue, and a network of water molecules. This aspect is expected to be crucial for the catalytic mechanism of protein kinases since charge stabilization at the TS is recognized to be a critical point for enzyme catalysis. In particular, the crystal structure of CDK2 used in this study contains the glycine-rich (Gly-rich) loop in the open state; this structural moiety functions as a lid closing the active site. It has been observed that the presence of two Mg^2+^ ions in the active site favors the closed state [21], where backbone amides of the Gly residues make hydrogen bonding interactions with the β-phosphate and presumably with the phosphoryl group at the TS. Thus, it is expected that in the presence of two Mg^2+^ ions the charge distribution on the phosphoryl group may be stabilized by specific hydrogen bonding and electrostatic interactions that favor the catalysis. However, how and how much these charge stabilizing interactions affect the phosphoryl transfer reaction is a question that remains open. These points are expected to be covered in future work.

## Conclusions

CDK2 is a very important kinase that is considered a study model for the cyclin-dependent kinase family. As other kinases, this enzyme catalyzes the phosphoryl transfer reaction from an ATP molecule to a peptide substrate containing a Ser o Thr residue to be phosphorylated. Though this system has been subject of experimental and computational studies, the two proposed reaction mechanisms, substrate-assisted and base-assisted, had not been studied until now within the same computational approach, which is a requisite in order to have a proper comparison between them. In this context, our results suggest that the base-assisted mechanism is the preferred path with an estimated potential energy barrier of 14.3 kcal/mol, which is in good agreement with the experimentally derived value. This mechanism is stepwise, in contrast to the substrate-assisted mechanism that is concerted. Both mechanisms show TSs that have a high dissociative character. Interestingly, a new feature in the base-assisted mechanism has been observed: a spontaneous proton transfer from Lys129 to one of the oxygen atoms of the transferred phosphoryl group. This event takes place late in the reaction progress, at the second TS, with the proposed effect of neutralizing the negative charge on the phosphoryl group.

Detailed mechanistic information has been given in terms of bond orders and natural population analysis. In both mechanisms, the breaking of the O_3β_-P_γ_ bond precedes the formation of the new bond with the entering oxygen from the substrate Ser residue. The rest of the bond breaking and bond forming events, related with the proton transfer to the phosphate group or to residue Asp127, occur in a synchronous fashion, although some degree of asynchronicity is observed for the base-assisted mechanism. Charge analysis showed that, due to the dissociative character of the mechanisms, the charge on the phosphoryl group becomes more positive when a metaphosphate species is formed. These results help to clarify how is the charge evolution on key atoms during the enzyme catalysis, such as the Mg^2+^ ion, the P_γ_ atom and the non-bridging oxygens of the phosphoryl group. Charge transfer effects appear to have an important role, specifically on the Mg^2+^ ion, which delivers electron density as the reaction moves to the product state. Also, it was observed that atomic charges on the non-bridging oxygens are modulated by local interactions from the protein microenvironment, mainly electrostatic and hydrogen bonding interactions with, primarily, the Mg^2+^ cofactor, active site residues such as Lys129, and water molecules. Future work will be concentrated in elucidating the effect of a second Mg^2+^ ion in the mechanism and its effect on the charge distribution of the phosphoryl group. In summary, these results help to advance towards a more comprehensive understanding of phosphoryl transfer reactions in protein kinases, information that can be highly relevant for the development of new inhibitors.

## Acknowledgments

The authors acknowledge to “Centro de Bioinformática y Simulación Molecular (CBSM)” at Universidad de Talca for providing computational resources to carry out the calculations reported in this study.

## Supporting information

**S1 Fig. Distance between the H_γ_ atom from the substrate Ser residue and the O_δ1_ atom from Asp127 showing a stable hydrogen bonding interaction during an unrestrained 2 ns MD simulation.**

**S1 Table. Calculated Wiberg bond orders, for the bonds that are broken and formed, for the substrate-assisted mechanism at the three stationary points.**

**S2 Table. Calculated Wiberg bond orders, for the bonds that are broken and formed, for the base-assisted mechanism at the five stationary points.**

**S3 Table. NPA charges (e) for the most relevant atoms and the PO_3_ group for both mechanisms at the different stationary points.**

